# The ameliorative effect of *C-Kit*^pos^ hepatic endothelial Mertk deficiency on nonalcoholic steatohepatitis

**DOI:** 10.1101/2024.08.08.607275

**Authors:** Seng-Wang Fu, Yu-Xuan Gao, Hui-Yi Li, Yi-Fan Ren, Jun-Cheng Wu, Zheng-Hong Li, Ming-Yi Xu

## Abstract

Recently, Mer tyrosine kinase (Mertk) and KIT proto-oncogene (C-Kit) were reported play a role in liver sinusoidal endothelial cells (LSECs) in patients with nonalcoholic steatohepatitis (NASH). In this study, lower levels of C-Kit and higher levels of Mertk/p-Mertk were confirmed in steatotic LSECs and in the livers of patients and mice with NASH. C-Kit was suggested to negatively regulate Mertk signaling in steatotic LSECs. The steatotic LSECs in which Mertk was knocked down displayed high fenestration and reduced expression of procapillarized CD31/VN; showed antiangiogenic features and decreased expression of proangiogenic VEGF/ERK1/2; and exhibited intact mitophagy and upregulation of the Pink1/Parkin pathway. Bone marrow transplantation (BMT) of *C-Kit*^pos^-BMCs^sh-Mertk^ to MCD mice could equivalently protect endothelial functions. Steatotic hepatocytes (HCs) or hepatic stellate cells (HSCs) cocultured with LSECs^sh-Mertk^ exhibited diminished lipid deposition; decreased expression of prolipogenic LXR/SREBP-1c, proinflammatory TNF-α/IL-6 and profibrotic α-SMA/ColI; and increased expression of prolipolytic FXR/ADPN. Similarly, the BMT of *C-Kit*^pos^-BMCs^sh-Mertk^ to MCD mice ameliorated NASH. *C-Kit*^pos^-LSECs that underwent Mertk cleavage were found to limit NASH progression. Therefore, Mertk deficiency should be a novel therapeutic agent for restoring LSECs in patients with NASH.

## Introduction

The obesity epidemic has resulted in a drastic increase in nonalcoholic liver disease (NAFLD) incidence. Nonalcoholic steatohepatitis (NASH) can progress to cirrhosis and hepatocellular carcinoma (HCC) and has become the leading and most rapidly growing cause of liver transplantation worldwide ^[1]^. However, the underlying mechanisms of this disease are still unclear.

Liver sinusoidal endothelial cells (LSECs) are located mainly in the sinusoidal lumen of liver sinusoids. LSEC dysfunction is considered critical for the progression to NASH, but restoring homeostasis is a promising approach for treating NASH ^[2]^. On the basis of our previous single-cell RNA-sequencing (scRNA-seq) study, we identified an important cluster of *C-Kit*^pos^ (KIT proto-oncogene, receptor tyrosine kinase^positive^)-LSECs involved in NASH progression ^[3]^. Duan et al. first reported that infusing *C-Kit*^pos^-LSECs into NASH mice could counteract age-associated senescence and steatosis by restoring the pericentral liver endothelium ^[4]^. However, the exact mechanism involving *C-Kit*^pos^-LSECs in NASH is still unclear.

We also found that C-Kit closely cross-talks with Gas6 (growth arrest-specific gene 6) signaling in these kinds of LSECs. Mertk (myeloid-epithelial-reproductive tyrosine kinase), a member of the Tyro-Axl-Mer (TAM) receptor tyrosine kinase family, is usually expressed at the highest level on macrophages and has ligands such as Gas6, which are free proteins that attach to apoptotic cells during efferocytosis ^[5]^. Roh et al. reported that tenofovir alafenamide can treat NASH by reducing the number of infiltrating *Mertk*^pos^ macrophages in mouse livers ^[6]^. From another perspective, Pastore et al. reported that *Mertk*^pos^ macrophages can modify the secretome to promote profibrotic features in hepatic stellate cells (HSCs) ^[7]^. However, the function of Mertk in LSECs in NASH has not been determined.

Herein, we showed that steatotic (as described for PA incubation) *C-Kit^neg^* (*C-Kit*^negative^)-LSECs express high levels of Mertk; promote a phenotype of capillarization and angiogenesis; inhibit mitophagy; and accelerate steatosis, inflammation and fibrosis in adjacent hepatocytes (HCs)/HSCs *in vitro*. In contrast, transplanting protective *C-Kit*^pos^-bone marrow cells (BMCs), especially those with Mertk knockdown, reversed NASH *in vivo*.

## Materials and Methods

### Human samples

A total of 6 individuals were involved. (1) Three patients with biopsy-confirmed severe NASH were enrolled, and the degrees of steatosis and fibrosis were based on pathological steatosis scores (F3: >50% steatosis) and Scheuer’s classification ^[8]^, with elevated serum ALT levels. (2) The control group consisted of 3 matched biopsy-confirmed AIH patients without histological NASH. The details of the cohorts are described in Table S1. All enrolled patients provided written informed consent, and the study was approved by the ethics committee of Shanghai East Hospital.

### Mouse model

Eight-week-old male C57BL/6 mice (procured from Shanghai Laboratory Animal Co. Ltd., Shanghai, China) were randomly assigned to 4 groups (each group, n=5): (1) the control group (chow diet for 8 weeks); (2) the MCD group (40% carbohydrate, 10% fat, deficient in methionine and choline for 8 weeks); (3) the MCD-based BMT-Neg group (MCD diet for 6 weeks and then transplantation of *C-Kit*^pos^-primary BMCs (pBMCs)^sh-NC^ for 2 weeks); and (4) the MCD-based BMT-Mertk(-) group (MCD diet for 6 weeks and subsequent transplantation of *C-Kit*^pos^-pBMCs^sh-Mertk^ for 2 weeks).

Donor pBMCs were first prepared. Second, magnetic activated cell sorting (MACS; Miltenyi Biotec, Auburn, CA, USA) was employed to obtain donor *C-Kit*^pos^-pBMCs. Third, sh-Mertk and sh-NC were separately transfected into *C-Kit*^pos^-pBMCs. Recipient mice underwent lethal irradiation before BMT. Prior to transplantation, donor cells (*C-Kit*^pos^-pBMCs^sh-Mertk^ and *C-Kit*^pos^-pBMCs^sh-NC^) were adjusted to a concentration of 5×10^7^ cells/ml for injection, and subsequently, 5×10^6^ cells were injected into the tail vein of each recipient mouse. All the BMT mice were sacrificed after 2 weeks ^[9]^. The animal study was approved by the Institutional Animal Care and Use Committee of Shanghai East Hospital.

### Histological staining

Mouse liver tissue sections were prepared and stained with H&E, Masson’s trichrome and ORO. Steatohepatitis, lipid droplets, fibrosis and inflammatory infiltration of liver were observed by light microscopy (Leica Microsystems, Wetzlar, Germany). Images were analyzed using Image J 1.8.0 software (National Institutes of Health, USA). A diagnosis of NASH activity score was evaluated by the pathologist according to the published criteria ^[10]^, which was determined by steatosis, inflammation and balloon swelling.

### Mouse primary cells isolation and culture

#### Primary LSECs

pLSECs were isolated from C57BL/6 male mice (standard diet as control group, MCD diet as MCD group for 6-weeks) as previously described ^[11]^. Mouse livers were perfused through the inferior vena cava with warm collagenase (Roche, Basel, Switzerland), and the portal vein was sectioned. The liver was removed and mechanically dissociated. Cells were isolated by centrifugation through Percoll gradients (Yeasen Biotech Co. Ltd.) to obtain pLSECs. The pLSECs were cultured in ECM media (ScienCell, Carlsbad, CA, USA) supplemented with 5% fetal bovine serum (FBS; Invitrogen, Carlsbad, CA, USA), antibiotics and antimycotics (Sigma-Aldrich, St. Louis, MO, USA).

#### Primary HCs, KCs and HSCs

Primary cells were isolated from C57BL/6 mice as our previously study ^[11]^. The pHCs and pKCs were isolated using a two-step collagenase digestion method and cultured with M199 medium (Gibco, Grand Island, NY, USA) supplemented with 10% FBS and antibiotics. The pHSCs were isolated using a gradient centrifugation method. After perfusing the livers with collagenase and pronase (Roche), the pHSCs were isolated by Nycodenz density gradient (Sigma-Aldrich) centrifugation. The cells were subsequently cultured in DMEM (HyClone South Logan, UT, USA) supplemented with 10% FBS and antibiotics. PHCs were isolated using a two-step collagenase digestion method. After 1-7 days in culture, the cells were harvested for the subsequent experiments.

#### Primary BMCs (pBMCs)

Isolation of pBMCs was referred to previously reported method ^[12]^. The 6-weeks C57BL/6 male mice (n=5) were euthanized utilizing CO_2_. Dissect the skin of bilateral lower limbs, isolate femur and tibia completely. Cut both ends of the tibia or femur, rinse the bone marrow cavity with buffer through a syringe. Filter the flush solution and centrifuge at 600G for 5 min (4℃) to obtain the resuspension of pBMCs. The pBMCs were subsequently cultured in DMEM supplemented with 10% FBS and antibiotics. Determine cell concentration with automated cell counter. Typical yield can range from 1×10^7^ to 1.2×10^7^ progenitor cells/mouse.

### MACS of *C-Kit*^pos^- and *C-Kit*^neg^-primary cells

Preparation of primary cells (pBMCs or pLSECs) was performed using MACS method (Miltenyi Biotec, Cologne, Germany). Pellets of cells were suspended in MACS buffer, and 1×10^7^ total cells were incubated with 20 μL of C-KIT microbeads for 15 minutes in a refrigerator (2-4°C). The LS column was washed with buffer and centrifuged to obtain *C-Kit*^neg^-cells. Remove the column from the separator and flush out the magnetically labeled cells with buffer to obtain *C-Kit*^pos^-cells.

### Cell line culture and treatment

TMNK-1 (human LSEC), HepG2 (human hepatoma cell) and LX2 (human HSC) were cultured in DMEM containing 10% FBS and antibiotics. Before *in vitro* experiment, cells were first incubated in serum-free media overnight. To induce a lipotoxic environment, TMNK-1 cells were pretreated with PA (200 μM; Sigma-Aldrich) or 3% BSA as the vehicle control for 24 hours. LX2 cells were pretreated with transforming growth factor-β1 (TGF-β1, 10 ng/ml; Sigma-Aldrich) for 24 hours.

### Cell transfection

For C-Kit knockdown, shRNAs targeting human C-Kit (5’-GCGACGAGATTAGGCTGTTAT-3’, also known as sh-C-Kit) or a nonsense sequence (such as sh-NC) were inserted into the pLent-U6-puro plasmid. For C-Kit overexpression, full-length human C-Kit (ACCESSION: NM_000222) was cloned and inserted into the pENTER plasmid (ov-C-Kit), and an empty vector (as a vector) was used as a control. All the vectors were purchased from Vigene Biosciences (Shandong, China). For Mertk knockdown, shRNAs targeting human Mertk (5’-GCTCAATCAGTGTACCTAATA-3’, also known as sh-Mertk) or sh-NC were inserted into the pLKO.1-puro plasmid. For Mertk overexpression, a full-length human Mertk (ACCESSION: NM_006343) was cloned and inserted into the pCMV-3×FLAG-SV40-Neo plasmid (ov-Mertk), and an empty vector (as a vector) was used as a control. Lipofectamine 3000 (Invitrogen, Carlsbad, CA, USA) was used for transfection. The effectiveness of the transduction procedure was investigated after 48 hours through qPCR.

### SEM

LSECs cultured on glass coverslips were fixed in 2.5% glutaraldehyde (Servicebio, Wuhan, China) buffered with poly-lysine buffer (pH7.4, Sigma-Aldrich) overnight at 4°C and post-fixed with 1% osmium tetroxide (Servicebio) in cacodylate buffer (pH7.4, Sigma-Aldrich) for 1 hour at 4°C. After dehydration in a graded series of ethanol solutions (Sinopharm, Shanghai, China), the cultured cells were dried in a critical point apparatus (K850, Quorum, Brighton, UK) and coated with gold in a vacuum coating unit (MC1000, HITACHI, Tokyo, Japan).

### EdU assay

Cell proliferation was evaluated by the EdU assay Kit (Beyotime). First, cells with different treatments were incubated with EdU solution for approximately 2 hours. Then, cells were fixed via 4% paraformaldehyde (Sinopharm), followed by permeabilization with 0.5% Triton X-100 (Beyotime) for approximately and 1% 4’,6-diamidino-2-phenylindole (DAPI, Beyotime) to distinguish the nuclei. Samples were viewed under a fluorescence microscope.

### Transwell Assay

After treatment, cells were collected for further cultivation in a transwell chamber (Corning). The upper chamber was coated with Matrigel (BD Bioscience, San Jose, CA, USA), and the lower chamber was filled with DMEM containing 10% FBS. Cells were cleared from the upper member with a sterilized cotton swab after incubation for 48 h, and then, cells in the lower chamber were fixed with 4% paraformaldehyde. The number of migrating cells was calculated under a light microscope after staining with 0.5% crystal violet solution (Sinopharm).

### Tube formation assay

After thawing, subculturing, and transfecting, the treated cells were digested with 0.25% trypsin (Procell, Wuhan, China) to create a single-cell suspension in serum-free medium. After counting, the cells were evenly seeded into 24-well plates precoated with Matrigel at a density of 1×10^5^ cells per well. After continuous culture for 8 hours, images were taken using an inverted microscope (Nikon, Tokyo, Japan).

### MtSOX and mtCMXRos

Mitochondrial ROS production was analyzed using a MitoSOX Red staining Kit (Invitrogen). The MMP was detected using a MitoTracker Red CMXRos Kit (Invitrogen). Representative images were captured using a TCS SP8 CARS fluorescence microscope (Leica Microsystems).

### Cell coculture

TMNK-1 cells were cocultured into 6 groups and incubated in the upper chambers. The cells were pretreated or transfected with BSA, PA, PA+sh-NC, PA+sh-Mertk, PA+vector or PA+ov-Mertk. Upper chamber cells were plated on polystyrene transwell plates (Corning, Inc.) with a 0.4 mm pore size at 3×10^5^ cells per well. Then, HepG2 or LX2 cells were plated in 6-well plates at 3×10^5^ cells per well. The upper cell-containing transwell was then placed into cell-containing 6-well dishes and cocultured for another 24 hours.

### PCR

qPCR was performed using a SYBR Green PCR Kit (Yeasen Biotech Co. Ltd., Shanghai, China) and an ABI 7900HT Fast Real-Time PCR System (Applied Biosystems, Foster City, CA, USA). The primer sequences (Sangon Biotech, Shanghai, China) are listed in Table S2.

### Western blot

Total protein was extracted from liver tissue or cells with a total protein extraction kit (KeyGEN, Nanjing, China), and the concentration was determined using a BCA protein concentration kit (GBCBIO, Guangzhou, China). Proteins were separated by 10% SDS‒PAGE and transferred to polyvinylidene fluoride membranes (Millipore, Billerica, MA, USA). After the membranes were blocked with 5% nonfat milk for 1 hour, they were incubated with the primary antibody at 4 °C overnight. The membranes were washed the next day and incubated with goat anti-rabbit HRP-conjugated IgG at 37 °C for 2 hours. The bands were detected using a chemiluminescent imaging system. The primary and secondary antibodies used are shown in Table S3.

### IF assay

Mouse liver slides or cell climbing slides were used for the IF assay. Antigen retrieval from the liver slides was performed using citrate buffer in a microwave. The cells on the slides were fixed with 4% paraformaldehyde for 15 minutes at 37 °C. Goat serum (5%; Boster, Wuhan, China) was used to block the samples for one hour. The slides were then incubated with primary antibodies overnight at 4 °C. DAPI was applied to visualize the nucleus. Representative images were captured via fluorescence microscopy. Relative fluorescence values were measured using ImageJ 1.8.0. The primary and secondary antibodies used are listed in Table S4.

### Statistical analysis

All the results shown are based on three repeated experiments. All the data are presented as the mean ± standard deviation. Differences between 2 groups were analyzed with an unpaired Student’s t test. All the statistical analyses were performed using SPSS 19.0 software (SPSS, Inc., Chicago, IL, USA). A *p* value <0.05 was considered to indicate statistical significance.

## Results

### C-Kit negatively regulates Gas6-Mertk signaling in LSECs in NASH

In our previous scRNA-seq study ^[3]^, a cluster of *C-Kit*^pos^-LSECs was first confirmed to mitigate NASH. Analysis of the differentially expressed genes (DEGs) and gene ontology (GO) biological functions revealed that C-Kit cross-talk with Gas6 was activated in *C-Kit*^pos^-LSECs (Table S5). Then, we analyzed the expression of Gas6 and TAM kinases in *C-Kit*^pos^- or *C-Kit*^neg^-pLSECs (primary LSECs) derived from wild-type (WT) mice. Similarly, the expression of C-Kit was lower, while that of Gas6 and Mertk/p-Mertk was greater in palmitic acid (PA)-treated pLSECs (Fig. 1A), *C-Kit*^neg^-pLSECs (Figs. 1A/B) and PA-treated TMNK-1 cells (Figs. 1E/F) than in control cells (*p*<0.05). However, the expression of Axl and Tyro3 did not differ among the groups of pLSECs (Fig. S1A).

**Figure 1.**
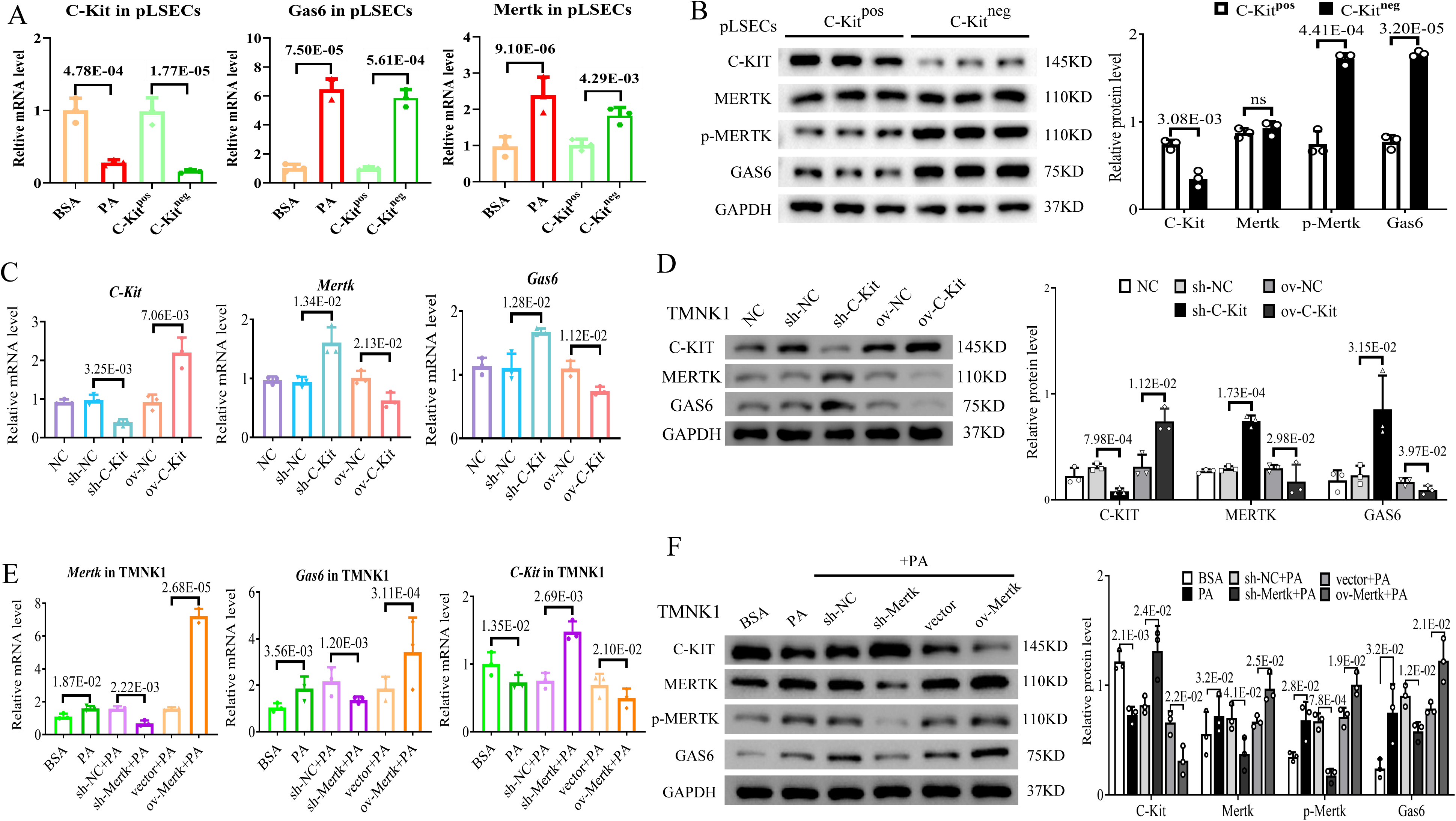
The regulatory relationship between C-Kit and Gas6-Mertk signaling in LSECs *in vitro*. The pLSECs were divided into 4 groups: BSA, PA, *C-Kit*^pos^ and *C-Kit*^neg^. The mRNA (A) and protein (B) levels of C-Kit, Gas6 and Mertk/p-Mertk were examined by qPCR and western blotting. TMNK-1 cells were divided into 5 groups: NC (blank control), sh-NC, sh-C-Kit, ov-NC and ov-C-Kit. The mRNA (C) and protein (D) levels of C-Kit, Gas6 and Mertk were examined. The TMNK-1 cells were divided into 6 groups: BSA, PA (200 μM), sh-NC+PA, sh-Mertk+PA, vector+PA and ov-Mertk+PA. The mRNA (E) and protein (F) levels of C-Kit, Gas6 and Mertk/p-Mertk were examined. All the data are expressed as the mean±SEM. Two-sample Student’s t tests were used for statistical analyses. *p* values indicate a significant difference compared to BSA/ *C-Kit*^pos^-LSECs, sh-NC/ ov-NC-transfected TMNK-1 cells, or BSA/ sh-NC+PA/ vector+PA-transfected TMNK-1 cells.

To determine whether C-Kit regulates Gas6-Mertk in LSECs, we transfected TMNK1 cells with a plasmid to knock down C-Kit (short hairpin RNA C-Kit, sh-C-Kit) or overexpress C-Kit (ov-C-Kit). Compared with those in control cells, the expression of Gas6-Mertk in sh-C-Kit cells was upregulated, whereas that in ov-C-Kit cells was downregulated (*p*<0.05; Figs. 1C/D). Otherwise, the effects of Mertk knockdown (sh-Mertk) or overexpression (ov-Mertk) on Gas6 and C-Kit were detected in steatotic TMNK1 cells (as PA treatment). The levels of C-Kit were greater and those of Gas6 were lower in steatotic sh-Mertk cells, while the opposite changes were observed in steatotic ov-Mertk cells versus their control cells (*p*<0.05; Figs. 1E/F). It is inferred that Mertk is a key factor negatively regulated by C-Kit in steatotic LSECs.

### Abundance of *C-Kit*^neg^-LSECs with Gas6-Mertk signaling activation in NASH

To determine the relationship between Gas6-Mertk signaling and *C-Kit*^pos^-LSECs in real-world NASH, we compared the manifestations of Gas6-Mertk signaling in severe NASH patients and MCD-induced NASH mice. The livers of autoimmunity hepatitis (AIH) patients and control mice contained abundant *C-Kit*^pos^-LSECs without MERTK proteins, but the livers of severe NASH patients and MCD mice contained massive numbers of *C-Kit*^neg^-LSECs with substantial MERTK proteins (Figs. 2A/C). Compared with those in the controls, the IF values in the MERTK/C-KIT area were nearly 6.7- and 6.9-fold greater in patients with NASH (Fig. 2B) and MCD mice (Fig. 2D), respectively. Compared with those in control mice, lower levels of C-Kit and higher levels of Gas6/Mertk/p-Mertk were detected in MCD mice (Figs. 2E/F). Then, the expression of C-Kit/Gas6-Mertk was investigated in primary hepatic cells. Gas6-Mertk was highly expressed in steatotic pLSECs/pKCs (primary KCs) and activated pHSCs (primary HSCs), while C-Kit was expressed at only low levels in steatotic pLSECs/pHCs (primary HCs, Fig. 2G). These findings suggested that C-Kit negatively regulates Mertk only in steatotic LSECs. Our data first revealed that activating Mertk is associated with an increase in *C-Kit*^neg^-LSECs in NASH.

**Figure 2.**
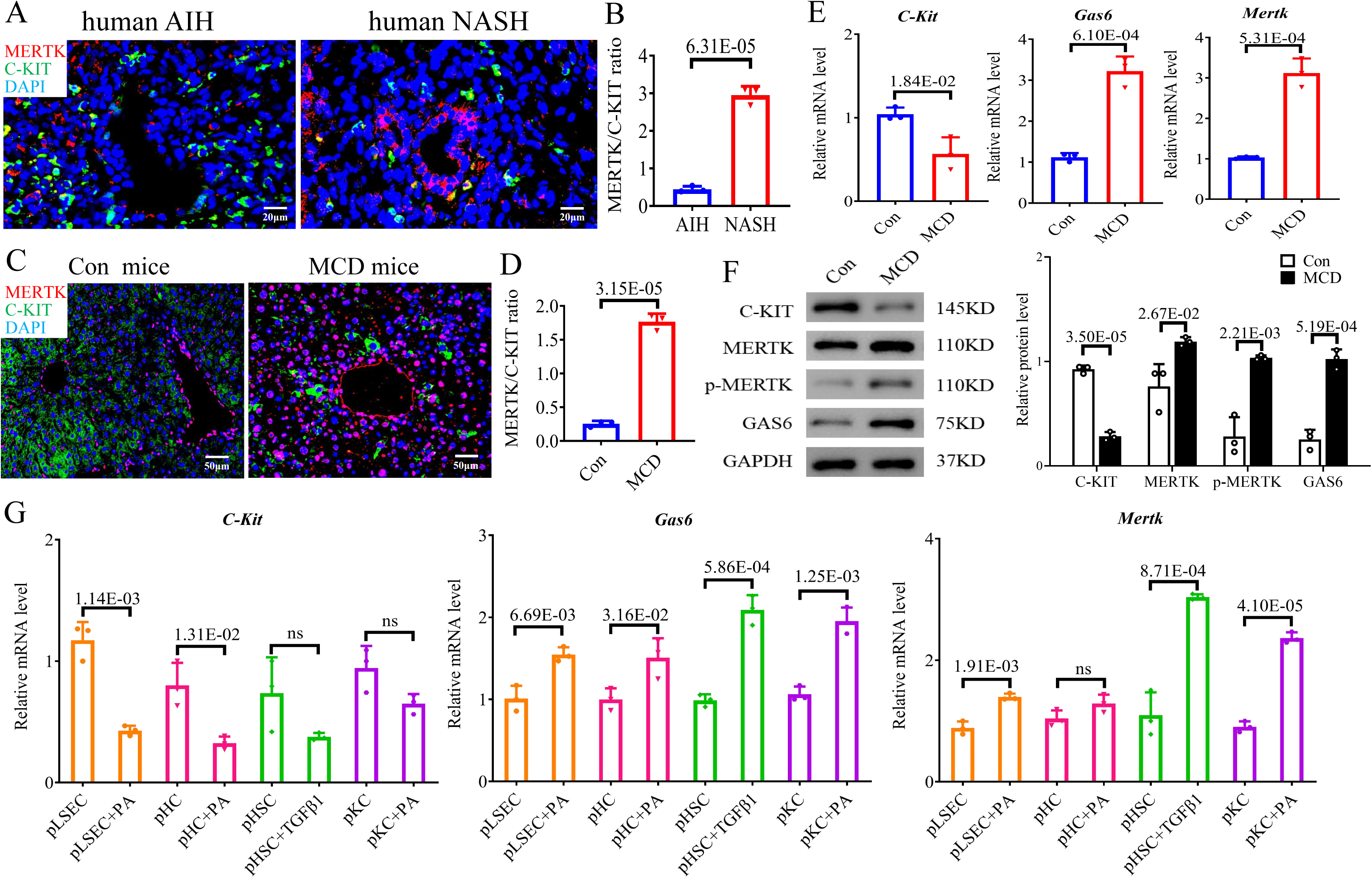
The expression of C-Kit and Mertk in NASH *in vivo and in vitro*. IF staining of MERTK (red)/C-KIT (green)/DAPI (blue) and quantification were shown for (A-B) human livers (AIH and NASH groups, each group n=3) and for (C-D) mouse livers (control and MCD groups, each group n=3). Scale bars=20-50 μm. The mRNA (E) and protein (F) levels of C-Kit, Gas6 and Mertk/p-Mertk were examined in the livers of the control and MCD groups. (G) The mRNA expression levels of C-Kit, Gas6 and Mertk were checked in the pLSEC±PA, pHC±PA, pHSC±TGF-β1, and pKC±PA groups. All the data are expressed as the mean±SEM. Two-sample Student’s t tests were used for statistical analyses. The *p* value indicates a significant difference compared to that of AIH patients, control mice, or pLSECs/ pHCs/ pHSCs/ pKCs.

### Mertk accelerates capillarization of LSECs in NASH

Capillarization, which is characterized by the defenestration of LSECs and the formation of a basement membrane in Disse, occurs in the early stage of NASH and impairs LSEC function as gatekeepers of liver homeostasis ^[13]^. To investigate whether Mertk promoted capillarization, 6 groups of differently expressed TMNK-1 cells [BSA, PA, sh-NC (nontargeting shRNA)+PA, sh-Mertk+PA, vector+PA, and ov-Mertk+PA] were studied. Through scanning electron microscopy (SEM), sinusoids of sh-Mertk+PA group contained more fenestrae clustered into sieve plates than control, while sinusoids of ov-Mertk+PA showed significant defenestrate (Fig. 3A). Quantitative morphometry showed that Mertk knockdown caused increase, while Mertk overexpression caused decrease in porosity (percentage of cell area covered by fenestration; Fig. 3B) and fenestration frequency (number of fenestrations per area; Fig. 3C) compared to those in the control groups (*p*<0.05). The continuous vascular endothelial markers CD31, CD34 and von Willebrand factor (VWF) are rarely expressed in LSECs; however, the increased expression of the above markers indicates sinusoid capillarization ^[14]^. Compared with those in the control group, the mRNA and protein levels of CD31/CD34 and VWF were greater in the PA group and ov-Mertk+PA group but lower in the sh-Mertk+PA group (*p*<0.05; Figs. 3D/E/S1B).

**Figure 3.**
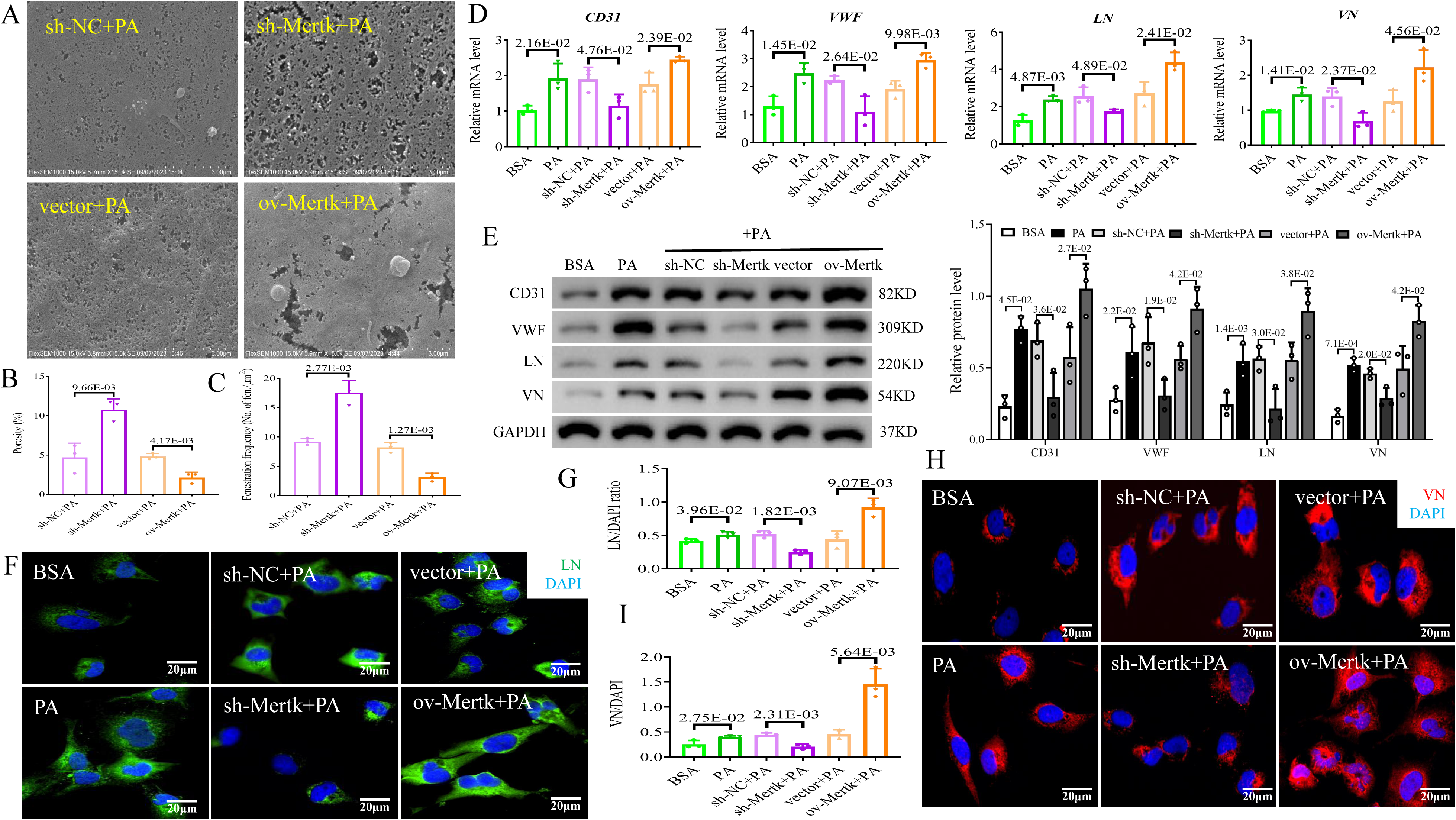
Mertk could facilitate capillarization of LSECs *in vitro*. Six groups of TMNK-1 cells were studied. (A) Representative SEM images of the 4 groups were shown (scale bars=3 μm). (B) The percentage of the cell area covered by fenestration and (C) the frequency of fenestration per area were quantified via SEM. The mRNA (D) and protein (E) expression levels of CD31, VWF, LN and VN were analyzed in the 6 groups via qPCR and western blotting. IF staining and quantification of (F-G) LNs (green)/DAPI (blue) and (H-I) VNs (red)/DAPI (blue) in the 6 groups (scale bars=20 μm). All the data are expressed as the mean±SEM. Two-sample Student’s t tests were used for statistical analyses. The *p* value indicates a significant difference compared to the BSA/ sh-NC+PA/ vector+PA group.

Laminin (LN) and vitronectin (VN) are major components of the basement membrane when excessively deposited in Disse, causing LSEC capillarization ^[15-16]^. Similarly, compared with those in the controls, the mRNA and protein expression levels of LN and VN in the PA and ov-Mertk+PA treatment groups were increased, but those in the sh-Mertk+PA treatment group were decreased (*p*<0.05; Figs. 3D/E). Immunofluorescence (IF) staining confirmed that the LN (green IF, Figs. F/G) and VN (red IF, Figs. H/I) proteins were indeed highly expressed in PA and ov-Mertk+PA cells but were obviously decreased in sh-Mertk+PA cells (*p*<0.05), suggesting that Mertk could promote endothelial capillarization in a steatotic environment. Since degradation of caveolin-1 (CAV-1) promotes LSECs defenestration ^[17]^, a profound mRNA reduction of CAV-1 was shown in PA and ov-Mertk+PA group, and increase of CAV-1 in sh-Mertk+PA group in contrast with each control group (*p*<0.05, Fig. S1B). In conclusion, Mertk can accelerate defenestration, increase basement membrane deposition, and ultimately promote the capillarization of LSECs in NASH.

### Mertk promotes the progression of LSECs into an angiogenic state in NASH

Adulteration leads to mild tissue hypoxia and the release of vascular endothelial growth factor (VEGF), which induces an angiogenic state in metabolic syndrome ^[18]^. To investigate the proangiogenic effect of Mertk on LSECs, a tube formation assay was performed. Compared with those in the corresponding controls, the vessel area, length and junctions in the PA and ov-Mertk+PA groups were greater, but the above vessel structures in the sh-Mertk+PA group were lessened (Fig. 4A). Quantitative morphometry revealed that PA and ov-Mertk+PA caused increase, while sh-Mertk+PA caused decrease in the total master segment length and the number of meshes compared to those in the control groups (*p*<0.05, Figs. 4B/C). VEGF and angiopoietin 1 (Ang1), which are angiogenic factors, promote blood vessel formation ^[19]^; VEGF can activate the downstream pathways of ERK, MAPK and Akt, which are involved in angiogenesis ^[20]^ during NASH. Moreover, VEGF, Ang1 and ERK/p-ERK1/2 mRNA and protein elevations were detected in the PA and ov-Mertk+PA groups, but their levels were lower in the sh-Mertk+PA group than in the control group (*p*<0.05; Figs. 4D/E). No differences in p38-MAPK or Akt mRNA levels were observed among the 6 groups (Figure S1C). IF staining of the VEGF protein also revealed a similar trend (red IF intensity of VEGF: PA vs. BSA, 2.17-fold; ov-Mertk+PA vs. vector+PA, 1.32-fold; sh-Mertk+PA vs. sh-NC+PA, 0.65-fold, *p*<0.05; Figs. 4F/G). The key LSEC dysfunctions involved in angiogenic enhancement and activation of VEGF/ERK signaling are associated with Mertk in NASH.

**Figure 4.**
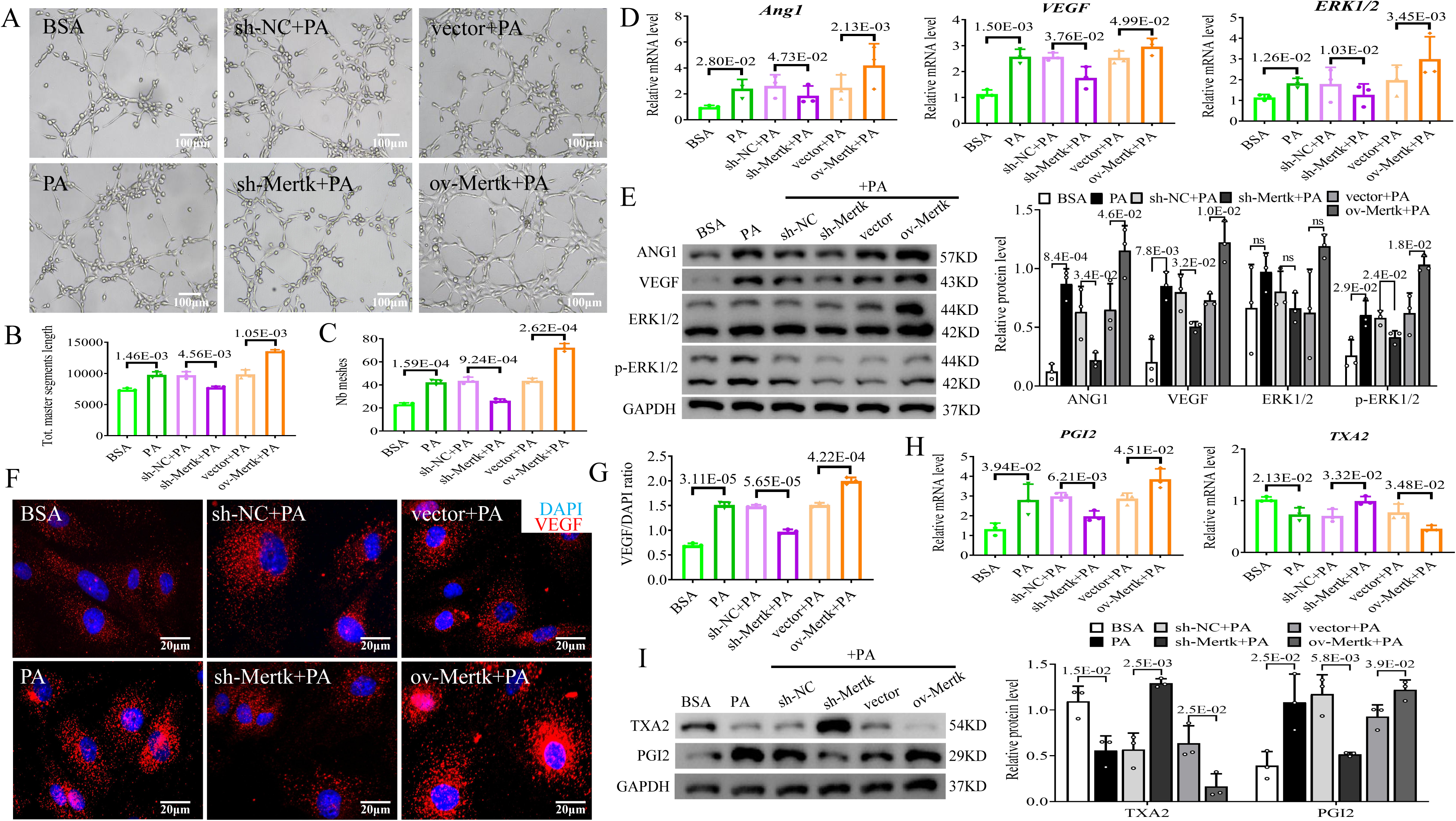
Mertk promoted angiogenesis and vasodilation of LSECs *in vitro*. Six groups of TMNK-1 cells were studied. (A) Representative images of tube formation assays in the 6 groups (scale bar=100 μm). (B-C) Quantitative analysis of tube formation, such as total master segment length and number of meshes, in the 6 groups. The mRNA (D) and protein (E) levels of Ang1, VEGF and ERK/p-ERK1/2 in the 6 groups were measured via qPCR and western blotting. (F-G) IF staining and quantification of VEGF (red)/DAPI (blue) in the 6 groups (scale bars=20 μm). The mRNA (H) and protein (I) levels of PGI2 and TXA2 in the 6 groups. All the data are expressed as the mean±SEM. Two-sample Student’s t tests were used for statistical analyses. The *p* value indicates a significant difference compared to the BSA/ sh-NC+PA/ vector+PA group.

### Mertk facilitates the proliferation, migration and vasodilation of LSECs in NASH

According to the results of the EdU incorporation assay, PA and ov-Mertk+PA treatment promoted the proliferation of LSECs, while sh-Mertk+PA treatment had the opposite effect compared to that of the controls (*p*<0.05; Figs. 5A/B). According to the results of the transwell assay, compared with those in the control group, the motility of LSECs in the PA or ov-Mertk+PA group was obviously increased, but their motility was apparently decreased in the sh-Mertk+PA group (*p*<0.05; Figs. 5C/D). NASH is accompanied by hepatic sinusoidal vasoconstriction/vasodilation dysfunction ^[21]^. The mRNA and protein levels of thromboxane A2 (TXA2, a vasoconstriction factor) were lower in the PA and ov-Mertk+PA groups but greater in the sh-Mertk+PA group than in the control group (*p*<0.05; Figs. 4H/I). Conversely, the mRNA and protein levels of prostaglandin-I-2 (PGI2, a vasodilatation factor) were greater in the PA and ov-Mertk+PA groups but lower in the sh-Mertk+PA group than in the control group (*p*<0.05; Figs. 4H/I). The mRNA levels of nitric oxide synthase (NOS, a vasodilation factor) and vascular cell adhesion molecule**-**1 (VCAM-1) were not significantly different according to PCR (Fig. S1D). In summary, Mertk stimulates LSEC proliferation and migration and enhances hepatic sinusoidal vasodilation in NASH patients.

**Figure 5.**
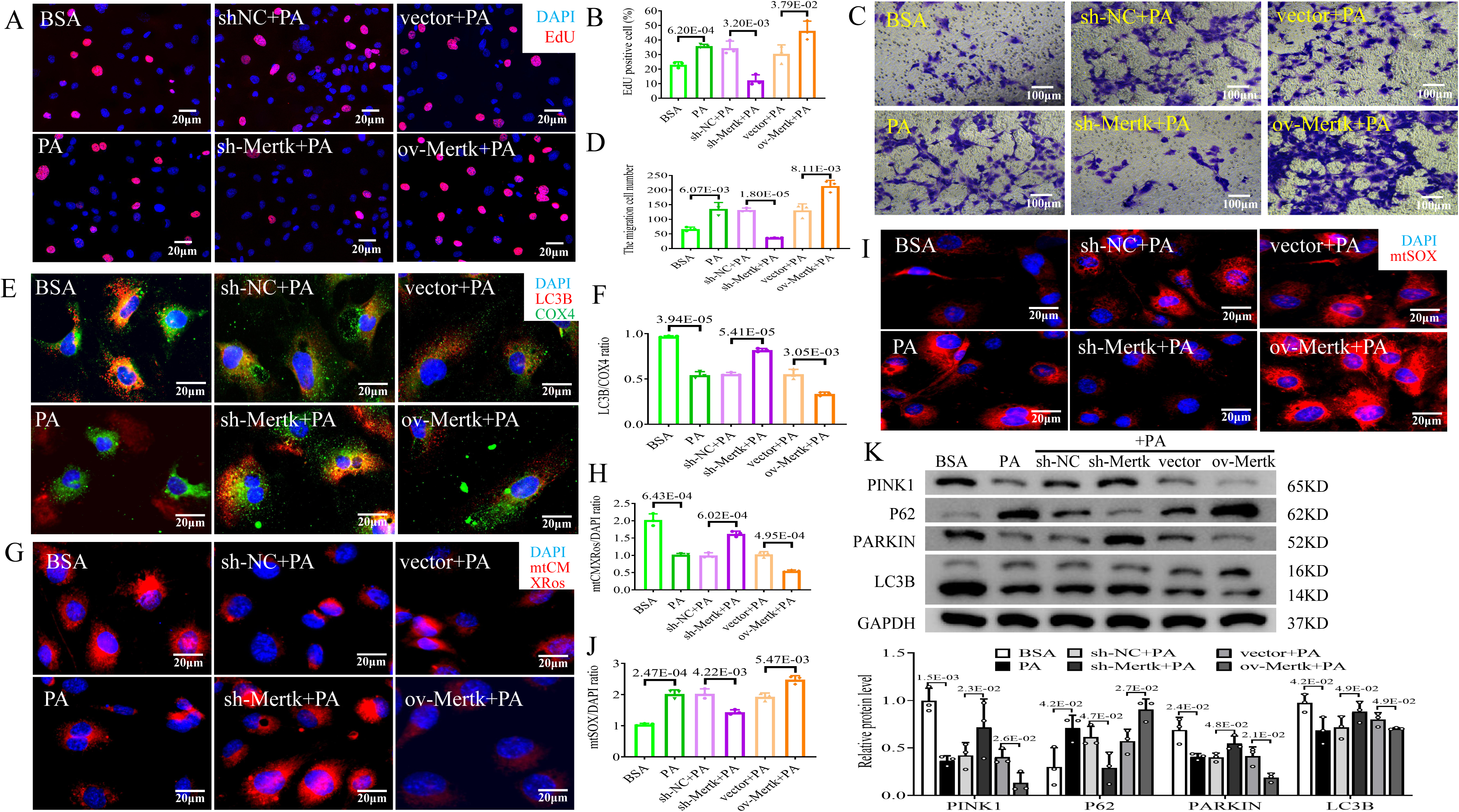
Mertk disrupted mitophagy but stimulated the proliferation and migration of LSECs *in vitro*. Six groups of TMNK-1 cells were studied. (A-B) Cell proliferation and quantitation were analyzed by EdU staining in the 6 groups (scale bar=20 μm). (C-D) Migration and migration were analyzed via transwell assays in the 6 groups (scale bar=100 μm). IF staining and quantification of (E-F) LC3B (red)/ COX4 (green)/ DAPI (blue); (G-H) mtCMXRos (red)/ DAPI (blue); and (I-J) mtSOX (red)/ DAPI (blue) in the 6 groups (scale bar=20 μm). (K) Protein levels of PINK1, P62, PARKIN and LC3B in the 6 groups determined via western blotting. All the data are expressed as the mean±SEM. Two-sample Student’s t tests were used for statistical analyses. The *p* value indicates a significant difference compared to the BSA/ sh-NC+PA/ vector+PA group.

### Mertk inhibits mitophagy in LSECs in NASH

Mitophagy, a mitochondrial quality control mechanism that removes dysfunctional mitochondria, is inhibited during the development of NASH ^[22]^. To evaluate the effect of Mertk on mitophagy in LSECs, costaining for light chain 3B (LC3B, a marker of autophagy; red IF) and cytochrome c oxidase subunit 4 (COX4, a marker of mitochondria; green IF) was performed via IF. Treatment with PA and ov-Mertk+PA decreased the colocalization of mitochondrial autophagy (orange IF), suggesting mitophagy deficiency; however, sh-Mertk+PA treatment increased this colocalization (0.56-fold decrease in the PA vs. BSA group; 0.61-fold decrease in ov-Mertk+PA vs. vector+PA group; 1.48-fold increase in sh-Mertk+PA vs. sh-NC+PA group; *p*<0.05; Figs. 5E/F). Mitophagy disturbance occurs along with the loss of the mitochondrial membrane potential (MMP) and excessive mitochondrial reactive oxygen species (ROS) production ^[23-24]^. PA and ov-Mertk+PA treatment could induce a decrease in the MMP, whereas sh-Mertk+PA treatment ameliorated mitochondrial depolarization, as shown by IF staining of mtCMXRos (MitoCMXRos) red (PA vs. BSA group: 0.50-fold; ov-Mertk+PA vs. vector+PA group: 0.54-fold; sh-Mertk+PA vs. sh-NC+PA group: 1.63-fold; *p*<0.05; Figs. 5G/H). Additionally, IF staining of mtSOX (MitoSOX) red revealed that PA and ov-Mertk+PA treatment could increase mitochondrial ROS production, which was alleviated by sh-Mertk+PA treatment (PA vs. BSA group: 1.94-fold; ov-Mertk+PA vs. vector+PA group: 1.28-fold; sh-Mertk+PA vs. sh-NC+PA group: 0.71-fold; *p*<0.05; Figs. 5I/J). The phosphatase and tensin homolog-induced putative kinase 1 (Pink1)/Parkin-dependent mitophagy signaling pathway has been shown to play an important role in the progression of NASH ^[25]^. We also detected proteins involved in the Pink1/Parkin axis in 6 groups of LSECs. The protein levels of PINK1, PARKIN and LC3B were significantly lower, and the protein level of P62 was greater in the PA and ov-Mertk+PA groups than in the control group, suggesting the inhibition of mitophagy (*p*<0.05; Fig. 5K). In contrast, sh-Mertk+PA treatment markedly rescued mitophagy by increasing PINK1, PARKIN, and LC3B levels and decreasing P62 levels compared with those in the control group (*p*<0.05; Fig. 5K). Taken together, these results suggest that Mertk inhibits the Pink1/Parkin-mediated mitophagy pathway in LSECs in NASH.

### Mertk knockdown in LSECs could alleviate NASH through inference of peripheral hepatic cells

Next, we assessed whether different Mertk expression in LSECs could directly regulate liver function by using a coculture system. HepG2 or LX2 cells were cocultured with 6 groups of TMNK-1 cells, namely, BSA, PA, PA+sh-NC, PA+sh-Mertk, PA+vector and PA+ov-Mertk. It is generally accepted that liver X receptor (LXR), sterol regulatory element binding protein-1c (SREBP-1c) and fatty acid synthase (FAS) promote lipogenesis, while farnesoid X receptor (FXR), adiponectin (ADPN) and peroxisome proliferator-activated receptor-α (PPAR-α) contribute to the progression of lipolysis. Lipid droplet accumulation increased (Figs. 6A/B), the levels of prolipogenic factors (LXR, SREBP-1c, and FAS) also increased, and the levels of prolipolytic factors (FXR, ADPN, and PPAR-α) decreased in HepG2 cells in the PA/PA+ov-Mertk group compared to those in the BSA/PA+vector group (*p*<0.05; Figs. 6C/E, S2A/C). These phenomena were reversed in the HepG2 cells in the PA+sh-Mertk group compared to those in the PA+sh-NC group (*p*<0.05; Figs. 6A/B/C/E, S2A/C).

**Figure 6.**
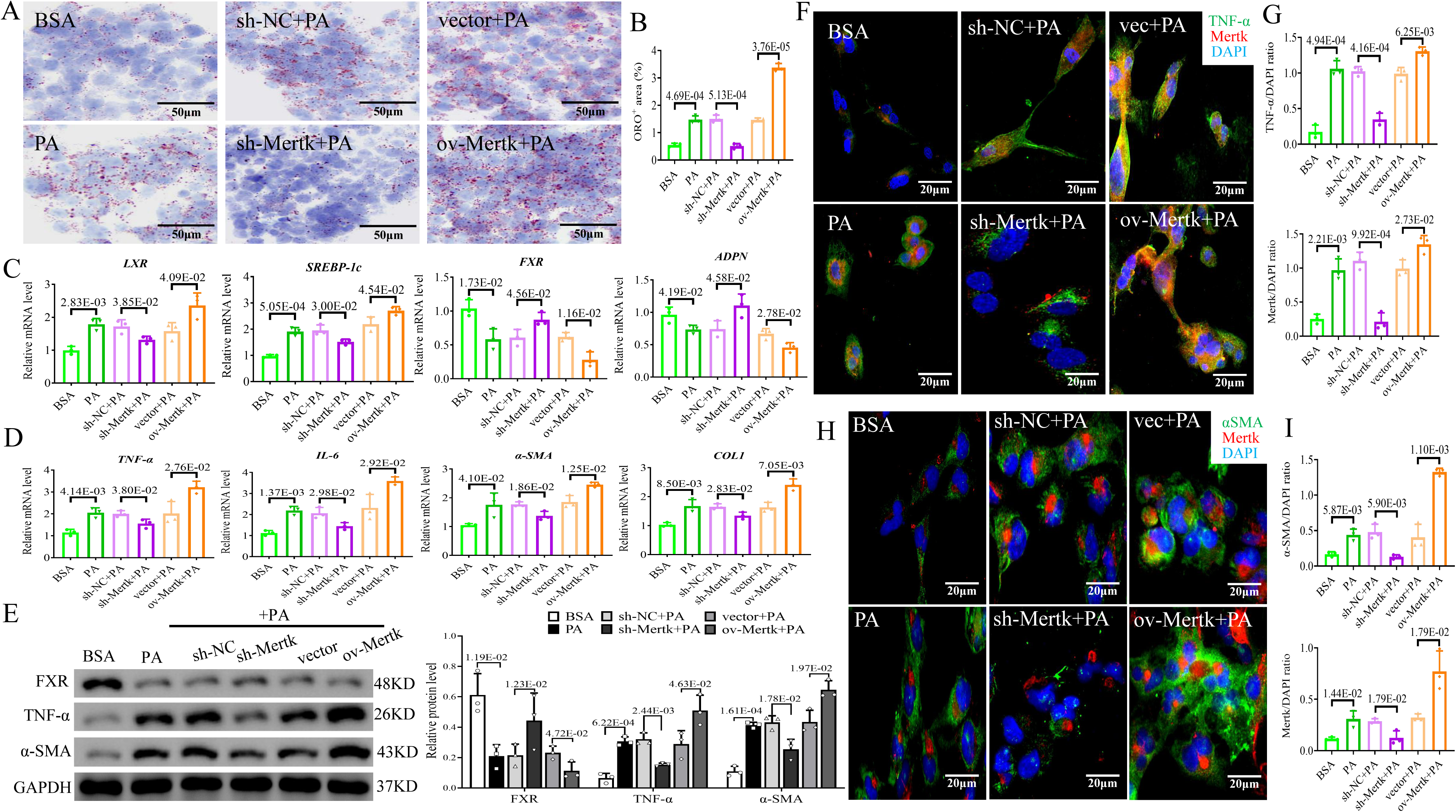
*In vitro* coculture experiments of LSECs and peripheral hepatic cells in NASH. HepG2 or LX2 cells were cocultured with 6 groups of TMNK-1 cells, namely, BSA, PA, PA+sh-NC, PA+sh-Mertk, PA+vector and PA+ov-Mertk. (A-B) ORO staining of HepG2 cells and quantification of the 6 groups (scale bar=50 μm). The mRNA levels of (C) LXR, SREBP-1c, FXR and ADPN in HepG2 cells; (D) TNF-α and IL-6 in HepG2 cells; and α-SMA and ColI in LX2 cells cocultured with 6 groups of TMNK-1 cells. (E) The protein levels of FXR, TNF-α and α-SMA in HepG2/LX2 cells cocultured with 6 groups of TMNK-1 cells. IF staining and quantification of (F-G) Mertk (red)/ TNF-α (green)/ DAPI (blue) in HepG2 cells; (H-I) Mertk (red)/ α-SMA (green)/ DAPI (blue) in LX2 cells cocultured with 6 groups of TMNK-1 cells (scale bar=20 μm). All the data are expressed as the mean±SEM. Two-sample Student’s t tests were used for statistical analyses. The *p* value indicates a significant difference compared to the BSA/ sh-NC+PA/ vector+PA coculture group.

In addition, the expression of proinflammatory factors, including tumor necrosis factor (TNF)-α and interleukin (IL)-6; IL-1β, in HepG2 cells; and profibrotic factors, including smooth muscle actin (α-SMA), collagen I (ColI), vimentin (Vim), and fibronectin (FN), in LX2 cells was upregulated in the PA/PA+ov-Mertk group compared to the BSA/PA+vector group (Figs. 6D/E, S2B/C). In contrast, the expression of these factors was downregulated in the HepG2/LX2 cells in the PA+sh-Mertk group compared to the PA+sh-NC group (Figs. 6D/E, S2B/C). IF costaining revealed increased levels of TNF-α (green IF, Figs. 6F/G) in HepG2 cells and increased levels of α-SMA (green IF, Figs. 6H/I) in LX2 cells, as did Mertk (red IF) in the PA/PA+ov-Mertk group compared to those in the control group. Additionally, the morphology of the PA+sh-Mertk group was greater than that of the PA+sh-NC group, as indicated by the decreased expression of TNF-α (Figs. 6F/G) and α-SMA (Figs. 6H/I) along with Mertk.

Subsequently, we explored the effect of Mertk on the proliferation of HCs cocultured with LSECs. EdU-positive cells were significantly greater (Figs. S3A/B), as were the mRNA and protein levels of CyclinD1 and PCNA (Figs. S3C/D), in HepG2 cells in the PA/PA+ov-Mertk group than in those in the BSA/PA+vector group. Additionally, the proliferation of the HepG2 cells in the PA+sh-Mertk group was diminished compared to that of the PA+sh-NC group (Fig. S3). These data indicate that Mertk knockdown in LSECs leads to a reduction in lipid accumulation, inflammation and fibrosis in hepatic cells *in vitro*.

### Bone marrow transplanting (BMT) of *C-Kit*^pos^-BMCs^sh-Mertk^ could alleviate NASH in mice

*In vitro* experiments confirmed that Mertk-mediated suppression of LSECs had a therapeutic effect on NASH. We further investigated the role of knocking down Mertk in *C-Kit*^pos^-BMCs through a BMT mouse model. *C-Kit*^pos^-BMCs transfected with sh-NC or sh-Mertk were separately transplanting into MCD-fed mice after lethal X-ray irradiation; thus named BMT-Neg and BMT-Mertk(-) groups, respectively. MCD-feeding led to the downregulation of C-Kit and upregulation of Gas6/Mertk/p-Mertk mRNA or proteins, as described above (Figs. S4A/B). After *C-Kit*^pos^-BMCs transplantation, the mRNA or protein expression of C-Kit was not differentially expressed; however, a noticeable decrease in Gas6/Mertk/p-Mertk was detected in BMT-Mertk(-) mice, indicating that the BMT-Mertk(-) model was successfully established (Figs. S4A/B).

Oil red O (ORO), hematoxylin-eosin (H&E) and Masson staining of the livers revealed that steatohepatitis and fibrosis were obviously exacerbated in the MCD diet-fed mice compared with those in the control group (Figs. 7A-D). However, transplanting *C-Kit*^pos^-BMCs^sh-Mertk^ markedly reduced hepatic lipidosis, lobular inflammation and collagen deposition compared to transplanting *C-Kit*^pos^-BMCs^sh-NC^ into MCD-fed BMT mice (Figs. 7A-D). An MCD diet caused an elevation in prolipogenetic LXR and SREBP-1c, but a decline in prolipolytic FXR and ADPN in MCD versus control group (Figs. 7E/G). As *C-Kit*^pos^-BMCs^sh-Mertk^ grafting, prolipogenetic factors were suppressed, while prolipolytic factors especially FXR was stimulated compared to *C-Kit*^pos^-BMCs^sh-NC^ grafting (Figs. 7E/G). Activators of inflammation (TNF-α and IL-6) and fibrosis (α-SMA and ColI) were also increased in the livers of MCD-fed mice compared with those of control mice, while *C-Kit*^pos^-BMCs^sh-Mertk^ grafting reversed these changes, particularly in the expression of TNF-α and α-SMA (Figs. 7F/G). To better characterize the correlation between Mertk and these genes, we compared the hepatic distribution of Mertk (red IF) and TNF-α (green IF) or α-SMA (green IF) by IF staining. Most of the Mertk signals associated with TNF-α- or α-SMA-positive cells were significantly greater in the MCD mice than in the control mice (Figs. 7H-J). Additionally, these proteins were deficient in the BMT-Mertk(-) group compared to the BMT-Neg group (Figs. 7H-J). Thus, BMT via *C-Kit*^pos^-BMCs^sh-Mertk^ to mice with NASH appears to reverse hepatic steatosis and fibrosis. This finding indicates that Mertk cleavage in LSECs limits NASH progression *in vivo*.

**Figure 7.**
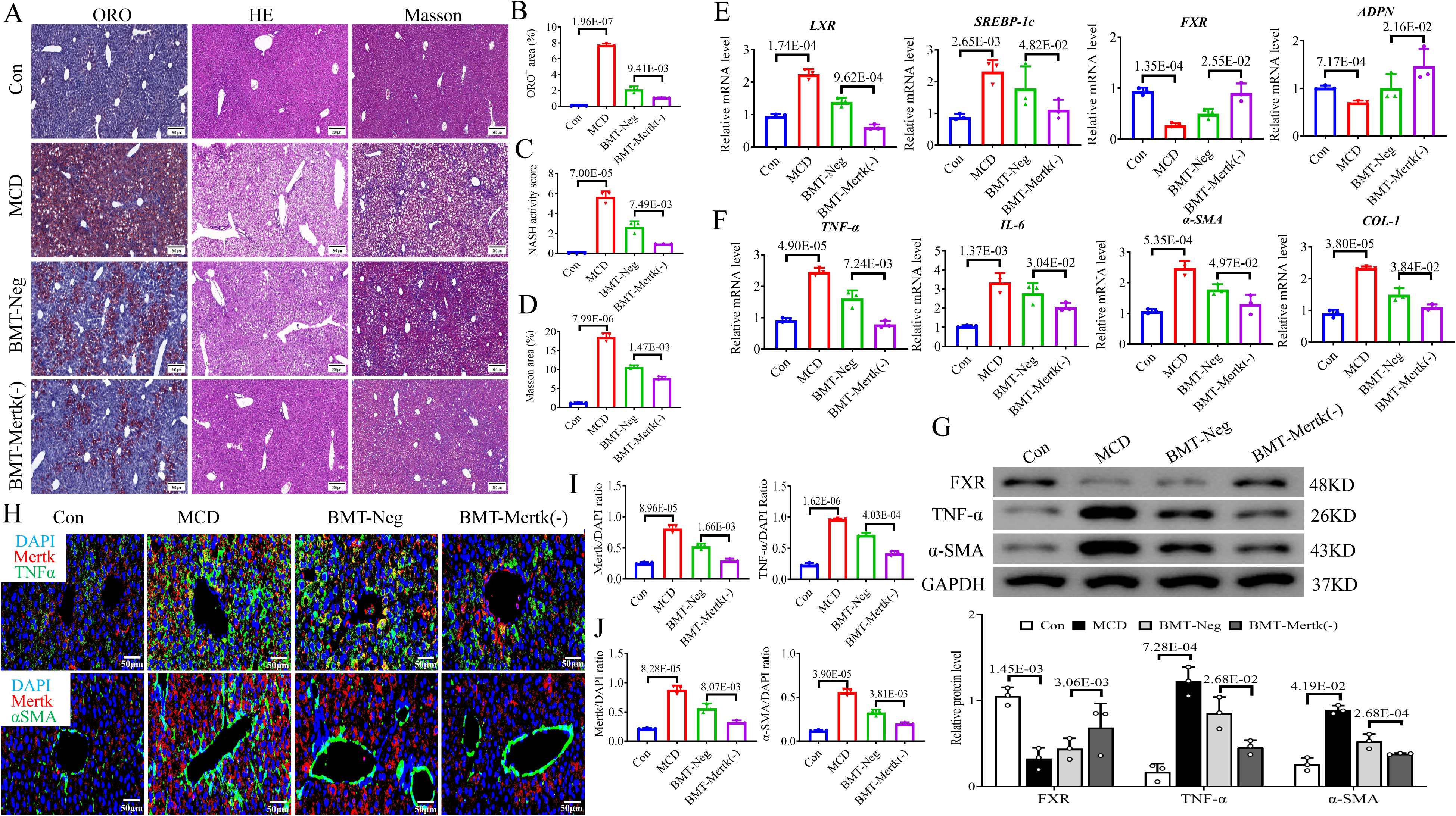
BMT of *C-Kit*^pos^-BMCs^sh-Mertk^ could alleviate NASH *in vivo*. The *in vivo* study was performed in 4 groups: Con (control-fed mice), MCD (MCD-fed mice), BMT-Neg (BMT of *C-Kit*^pos^-BMCs^sh-NC^ to MCD-fed mice), and BMT-Mertk(-) (BMT of *C-Kit*^pos^-BMCs^sh-Mertk^ to MCD-fed mice). (A-D) Representative images and quantitative analysis of ORO, H&E, and Masson staining of livers from the 4 groups (scale bar=200 μm). The mRNA expression levels of (E) LXR, SREBP-1c, FXR and ADPN and (F) TNF-α, IL-6, α-SMA and ColI in the 4 groups were measured via qPCR. (G) The protein levels of FXR, TNF-α and α-SMA in the 4 groups were measured via western blotting. (H-J) IF staining images and quantitative analysis of Mertk (red), TNF-α (green), DAPI (blue), and Mertk (red), as well as α-SMA (green) and DAPI (blue), in liver sections from the 4 groups (scale bar=50 μm). All the data are expressed as the mean±SEM. Two-sample Student’s t tests were used for statistical analyses. The *p* value indicates a significant difference compared to the Con/ BMT-Neg group.

### *C-Kit*^pos^-BMCs^sh-Mertk^ BMT also restored hepatic endothelial functions in NASH mice

Similarly, compared with those in control mice, the mRNA and protein expression levels of procapillarized factors (CD31, CD34, VWF, LN and VN) and proangiogenic factors (VEGF and Ang1) were significantly greater, while the incapillarized factor (CAV-1) was lower in MCD-fed mice (Figs. 8A-B/S4C-E). BMT from *C-Kit*^pos^-BMCs^sh-Mertk^ effectively inhibited the expression of procapillarized CD31 and VN and proangiogenic VEGF compared to BMT from *C-Kit*^pos^-BMCs^sh-NC^ to MCD-fed mice; other factors were not involved (Figs. 8A-B/S4C-E). CD31, VN and VEGF are commonly used as sinusoid capillarization and angiogenesis markers, and they are scattered throughout the normal liver. We detected CD31 (green IF), VN (red IF) and VEGF (red IF) expression in the majority of LSECs in MCD-fed livers but low expression in normal livers by IF staining (Figs. 8C-F). In contrast to the BMT-Neg group, which exhibited high expression of these 3 proteins, the BMT-Mertk(-) group exhibited low expression in LSECs (Figs. 8C-F). We further assessed mitophagy flux in MCD-BMT mice treated with different BMCs. IF staining for COX4 (green IF) and LC3B (red IF) revealed that the number of cells with colocalization of orange among the liver parenchyma and portal area was significantly lower in the MCD-fed mice than in the control mice (*p*<0.01; Figs. 8G-I). Furthermore, BMT-Mertk(-) mice had brighter orange IFs in those areas than did BMT-Neg mice (*p*<0.01; Figs. 8G-I). Stimulation of the Pink1/Parkin-mediated signaling pathway exacerbates mitophagy. A decreasing mRNA and proteins expressions of Pink1, Parkin and LC3B, and increasing expressions of p62 were found in MCD versus control mice (Figs. 8J/K). The BMT of *C-Kit*^pos^-BMCs^sh-Mertk^ exhibited an increase in Pink1/Parkin pathway-related factors compared to the BMT of *C-Kit*^pos^-BMCs^sh-NC^ (Figs. 8J/K). These results suggest that inhibiting Mertk in *C-Kit*^pos^-BMCs could effectively resuscitate Pink1/Parkin-mediated mitophagy in NASH *in vivo*.

**Figure 8.**
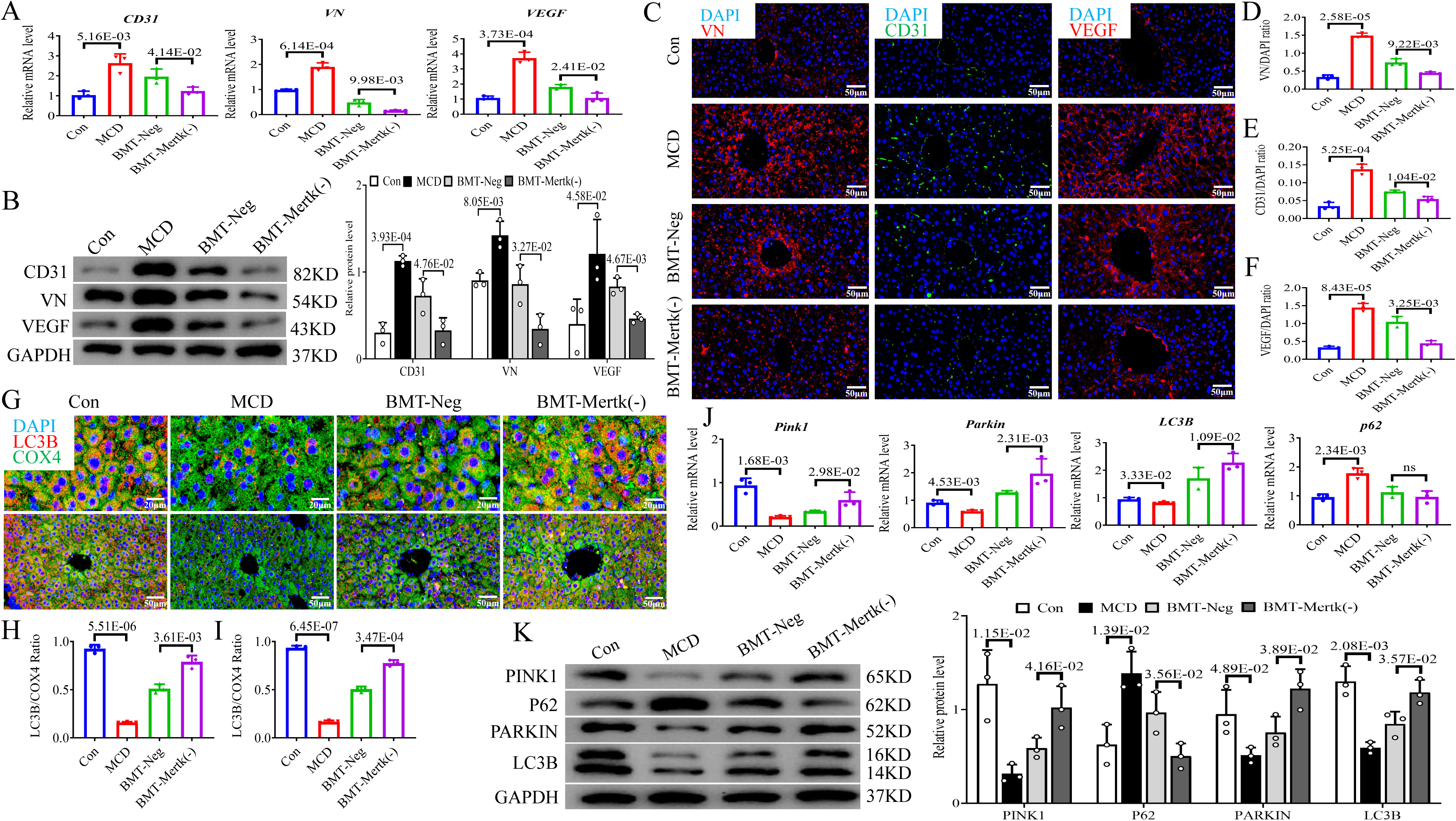
BMT of *C-Kit*^pos^-BMCs^sh-Mertk^ could ameliorate endothelial function in NASH *in vivo*. The *in vivo* study was performed in 4 groups. The mRNA (A) and protein (B) expression levels of CD31, VN and VEGF in the 4 groups were determined via qPCR and western blotting. (C-F) IF staining images and quantitative analysis of LN (red)/ DAPI (blue), CD31 (green)/ DAPI (blue), and VEGF (red)/ DAPI (blue) in the 4 groups (scale bar=50 μm). (G-I) IF staining images and quantitative analysis of LC3B (red), COX4 (green), and DAPI (blue) in the parenchymal and portal areas of the livers in the 4 groups (scale bar =50 μm). The mRNA (J) and protein (K) expression levels of Pink1, Parkin, LC3B and p62 in the 4 groups. All the data are expressed as the mean±SEM. Two-sample Student’s t tests were used for statistical analyses. The *p* value indicates a significant difference compared to the Con/ BMT-Neg group.

## Discussion

NAFLD is the most common liver disease and is characterized by hepatic steatosis ^[26]^. NASH patients with both cirrhosis and cardiovascular complications have the highest risk of mortality and morbidity ^[27]^. The US FDA has not approved a specific drug for NASH treatment; therefore, developing a new therapeutic drug is essential ^[28]^.

Endothelial cells (ECs) form interjunctions and regulate molecules and cells between the circulation blood and tissues. LSECs are characterized by the presence of transcellular pores as fenestrae and exhibit anti-inflammatory and antifibrotic features under physiological conditions ^[2]^. Recent scRNA-seq studies have defined several distinct subpopulations of LSECs with different gene expression profiles and elucidated their mechanism in NASH ^[29]^. Our previous scRNA-seq analysis revealed that a cluster of *C-Kit*^pos^-LSECs has a therapeutic effect on NASH^[3]^. Additionally, Duan et al. demonstrated that C-Kit deletion can perturb LSEC homeostasis and worsen hepatic steatosis, fibrosis and inflammation in NASH mice ^[4]^. However, the mechanism of action of *C-Kit*^pos^-LSECs in NASH has not been completely studied.

In the present study, bioinformatics analysis revealed that C-Kit in steatotic LSECs cross-talks with Gas6/TAM signaling to exercise biological functions. C-Kit expression is low, but Gas6-Mertk expression is high in steatotic LSECs (primary/cell line) and *C-Kit*^neg^-pLSECs. Further *in vitro* transfection experiments confirmed that C-Kit and Gas6-Mertk have mutual negative regulatory effects on steatotic LSECs. Coincidentally, NASH-triggered *C-Kit*^neg^-LSECs accumulate along with multiple Mertks in the sinusoidal portal area in humans and mice. Finally, there is no relevant C-Kit or Gas6-Mertk literature available, and there is also no consensus on this topic in NASH patients.

Mertk, a member of the TAM kinase family, is expressed in many tissues and is correlated with various cellular processes ^[30-31]^. Interestingly, all TAM kinases participate in the regulation of endothelial permeability. Mertk is also expressed in ECs ^[32]^ and is implicated in the maintenance of the blood-brain barrier ^[33]^. Li et al. reported that Mertk knockdown could alter adherens junction structure and decrease junction-related protein levels and basal Rac1 activity in ECs ^[34]^. Happonen et al. demonstrated that Mertk is a central regulator of angiogenesis, blood-brain barrier integrity, and blood coagulation in the mature vasculature ^[35]^. Only *Mertk*^pos^ macrophages have been confirmed to accelerate NASH-related inflammation and fibrosis ^[6-7]^, while the mechanism through which Mertk affects LSECs in NASH patients has not been determined.

We identified a new signaling pathway controlled by Mertk in *C-Kit*^pos^-LSECs involving three key aspects of NASH-related endotheliopathy. The first is capillarization of LSECs, which occurs early during NAFLD pathogenesis and in turn promotes hepatic steatosis. Using SEM, Miyao et al. observed sinusoidal capillarization in mice as early as one week after feeding on an MCD diet ^[36]^. Herrnberger et al. speculated that lost fenestrae and chylomicron remnants in mice are trapped in the circulation and no longer reach HCs, resulting in hepatic steatosis ^[37]^. According to our comprehensive SEM, IF staining and PCR/WB analyses, Mertk overexpression caused a decrease in fenestration porosity and frequency; multitudinous staining of LN/VN proteins; and increased in the mRNA and protein levels of capillarization-related CD31/CD34, VWF, LN, and VN in steatotic LSECs. However, Mertk knockdown had the opposite effect. *In vivo*, BMT of *C-Kit*^pos^-BMCs^sh-Mertk^ effectively inhibited hepatic CD31 and VN mRNA and protein expression in the livers of NASH mice. Thus, Mertk promotes the capillarization of LSECs in NASH, and inhibiting Mertk may reverse this change. The second is that LSECs acquire angiogenetic properties as NASH progresses to an advanced stage ^[38]^. Moreover, Mertk can regulate the expression of EC genes that encode regulators of angiogenesis ^[35]^. In our study, Mertk overexpression promoted angiogenesis by increasing vessel area, length and junctions and elevating the expression of proangiogenic VEGF, Ang1 and ERK/p-ERK1/2 in steatotic LSECs. Our data were again obtained via Mertk-promoted phosphorylation of ERK1/2, and this general pathway also operates for VEGF receptor signaling systems in ECs ^[39]^. Even so, specific inactivation of Mertk in steatotic LSECs results in antiangiogenic effects. When *C-Kit*^pos^-BMCs^sh-Mertk^ cells were subjected to BMT, the mRNA and protein expression of VEGF were also decreased in the livers of NASH mice. Activation of angiogenic VEGF-ERK1/2 signaling in LSECs is associated with Mertk in NASH; however, Mertk deficiency could reverse this effect. The third dysfunction in our study was an increase in proliferation and migration and the high expression of the vasodilation-related gene PGI2 when Mertk was overexpressed in steatotic LSECs. Additionally, Mertk deletion reversed these changes. Kumei et al. demonstrated that the PGI2-IP system plays a suppressive role in the progression of NASH ^[40]^, which contradicts our findings. Mertk stimulates proliferation and migration and promotes hepatic sinusoidal vasodilation in NASH; however, Mertk ablation ameliorates these dysfunctions.

However, mitochondria are key regulators of EC homeostasis by controlling signaling responses to the environment ^[41]^. It has been demonstrated that reducing mitochondria-derived ROS (mtROS) can ameliorate hepatic steatosis, substantiating the crucial role of mtROS in EC dysfunction ^[42-43]^. Hyperglycemia or hyperlipidemia cause mitochondrial dysfunction, including opening of permeability transition pores and release of proapoptotic cytokines, leading to mitochondrial and EC apoptosis ^[44-45]^. In our study, IF staining of COX4/LC3B, mtCMXRos and mtSOX revealed mitophagy and MMP damage but excessive mtROS production in steatotic LSECs overexpressing Mertk, especially those in the Pink1/Parkin pathway, was inhibited. Interestingly, Mertk suppression markedly rescued mitophagy through stimulating the Pink1/Parkin pathway in steatotic LSECs both *in vitro* and *in vivo*. Collectively, these studies support the vital role of mitochondrial impairment in NASH endotheliopathy. We hypothesize that Mertk knockdown in LSECs will improve mitophagy through the Pink1/Parkin pathway in NASH.

In addition to contributing to the progression of NASH, LSECs may respond to the release of proinflammatory activators, including TNF-α or IL-6/1β ^[46]^, secrete several chemokines and stimulate leukocyte recruitment into the liver parenchyma ^[47]^. LSECs play a major role in maintaining HSC quiescence via paracrine signaling ^[48]^. Capillarized LSECs accompanied by loss of fenestrae accelerate HSC activation and subsequent fibrosis ^[49]^. Importantly, we identified the interaction between LSECs and HCs/HSCs using an *in vitro* coculture system. Lipid droplet accumulation, increased levels of prolipogenic LXR/SREBP-1c/FAS, and decreased levels of prolipolytic FXR/ADPN/PPAR-α were detected in HCs cocultured with steatotic LSECs overexpressing Mertk. Then, proinflammatory factors (TNF-α and IL-6/1β) and profibrotic factors (α-SMA, ColI, Vim, and FN) were stimulated in HCs/HSCs cocultured with steatotic LSECs overexpressing Mertk. These phenomena could be favorable in HCs/HSCs cocultured with steatotic LSECs from Mertk knockdown mice. We further checked the therapeutic effect of BMT on *C-Kit*^pos^-BMCs^sh-Mertk^ in a NASH mouse model. When *C-Kit*^pos^-BMCs^sh-Mertk^ was used to treat NASH mice, hepatic lipidosis, lobular inflammation and collagen deposition were ameliorated, prolipogenetic factors (LXR and SREBP-1c) were downregulated, and prolipolytic factors (FXR and ADPN) were upregulated; moreover, proinflammatory factors (TNF-α and IL-6) and profibrotic factors (α-SMA and ColI) were also inhibited in the liver. Therefore, Mertk knockdown in LSECs led to a reduction in lipid accumulation, inflammation and fibrosis in hepatic cells both *in vitro and in vivo*. These findings indicate that LSECs capable of Mertk cleavage limit NASH progression.

Finally, our study similarly reveals a positive feedback loop between *C-Kit*^neg^-LSEC-related endotheliopathy and NASH progression. However, we first suggest that Mertk knockdown has a therapeutic effect on NASH in *C-Kit*^pos^-LSECs. Hence, restoring LSECs during NASH is a promising future treatment.

## Data Availability Statement

The datasets generated during and/or analyzed during the current study are available from the corresponding author on reasonable request.

## Acknowledgements

This study was supported by the National Natural Science Foundation of China (No. 82270604, 82100607, 82200629), Shanghai East hospital talent introduction project (No. DFRC2022005), Medical discipline Construction Project of Pudong Health Committee of Shanghai (No. PWYgf2021-02).

The funders had no role in the study design, data collection and analysis, decision to publish, or preparation of the manuscript.

## Author Contributions

Conceived the study design: M.Y.X. and Z.H.L.. Performed the experiments and analysed the data: S.W.F., Y.X.G., H.Y.L., Y.F.R. and J.C.W.. Wrote the manuscript: M.Y.X., S.W.F., Y.X.G. and Z.H.L.

